# Mesophilic compostability of polylactic acid and the associated microbiome as revealed by metagenomics

**DOI:** 10.1101/2025.06.29.662191

**Authors:** Shu Wei Hsueh, Anya Callista Kurniadi, Tan S. M. Amelia, Chin-Fa Lee, Sebastian D. Fugmann, Shu Yuan Yang

## Abstract

Polylactic acid (PLA), the most popular bioplastic, has high sustainability potential as it is bio-sourced and also harbors biodegradability. A form of its biodegradability is via composting, and it was previously established that thermophilic temperatures are needed for PLA breakdown in composts. Here we report the development of composts that have overcome the temperature requirement needed for PLA composting. Our mesophilic composts exhibited clear PLA biodegradability, and this is due to specific biological activity enriched in our material. To investigate the nature of this mesophilic activity, we conducted metagenomics analysis to reveal the microbial composition and enzyme-coding potential associated with PLA biodegradation. These efforts revealed multiple enzyme subtypes with strong enrichment on PLA surfaces in our trained composts, and the top candidate was a type of hydro-lyase, an enzyme that can cleave carbon-carbon and carbon-oxygen bonds, both present in the chemical structure of PLA, in the absence of water. Hydro-lyases represent a novel class of enzymes that could facilitate PLA degradation, and our results point to the model that the combinatorial action of multiple types of enzymes is what drives PLA biodegradation and how the temperature barrier for PLA composting is overcome.

## Introduction

Degradation of the bio-sourced polylactic acid (PLA) polymer could be biotic or abiotic, and biodegradation is thought to require less energy and produce fewer toxic byproducts, thereby being more environmentally favorable. The best-known form of PLA biodegradation is composting at industrial scales with internal reaction temperatures reaching 55-60°C and above [1]. Under such thermophilic conditions, PLA can be broken down first by surface hydrolysis which subsequently enables enzymatic depolymerization [2]. In composts of smaller volume that can only reach mesophilic temperatures, PLA degradation rates are minimal, presumably due to the lack of chemical hydrolysis [3].

Given the hydrophobicity and inertness of plastic surfaces that would be incompatible with the hydrophilic nature of enzymatic reactions, two-step models have been invoked to explain biodegradation of plastics, including PLA, in which biotic breakdown takes place after surface digestion or functionalization which could be biotic or abiotic [4]. In thermophilic composts, abiotic hydrolysis would achieve surface functionalization of PLA followed by further breakdown by microbial enzymes [2,5,6]. The microbes that contribute to PLA compostability are complex, thus it is challenging to tease apart individual contributions within compost systems. In comparison, studies using defined strains or enzymes have presented clearer pictures of how PLA depolymerization could occur. A variety of microbes including bacteria and fungi have been reported to exhibit PLA depolymerization activity [7,8], and among them, the filamentous Actinomycetes are well represented [9]. Nonetheless, there does not appear to be species specificity for PLA biodegradability.

The majority of enzymes that have been shown to act as PLA depolymerases are hydrolases including serine proteases, lipases, and esterases, and proteinase K is currently the benchmark example [10]. The reaction conditions performed have varied considerably with regards to temperature and substrate type, but multiple enzymes have been reported to be active in the mesophilic temperature range. This is interesting considering the requirement of PLA composting at elevated temperatures, and one possible explanation is that if PLA substrates used were of lower molecular weight, depolymerizing enzymes could more easily hydrolyze and access polymers in the absence of abiotic surface functionalization [11]. However, several studies reported depolymerization activity with high molecular weight PLA which is what is used for commercial products [12–14]. This is suggestive of the potential of biotic mechanisms in bypassing the thermophilic requirement of PLA biodegradation.

Here, we report our success in cultivating mesophilic composts with clear PLA biodegradability. Their emergence strongly suggested that microbial activities operating at lower temperatures can be sufficient for PLA to undergo biodegradation. To investigate the microbial strains and enzymes underlying such novel activity, we performed metagenomics sequencing of the microbiome on the surface of PLA material from our PLA-degrading mesophilic composts.

## Materials and methods

### Mesophilic composts and tests of PLA biodegradability

Our trained mesophilic composts that exhibited PLA biodegradability (thereafter referred to as trained compost) were cultivated through continuous exposure to PLA material. For their maintenance there was a consistent presence of commercially-purchased PLA products with 100% compostability certification around 0.1% volume in the compost bins, and the bins were otherwise fed with standard green waste mixes consisting of 75% raw fruit and vegetable waste, 12% garden waste, 6.5% wood dust, and 6.5% coffee grounds. This standard green waste was also what was used as untrained compost, the term used for composts that did not exhibit PLA biodegradability. For tests of PLA biodegradability of different trained compost bins, a fresh bin of untrained compost was set up for 14 days before mixing with trained material from specific bins at a 1:1 volume ratio. The mixtures were verified for general compostability by monitoring two pieces of materials with high degradability, a piece of paper bag (100 μm) and a piece of paper kitchen towel (70 μm), both 2.5×2.5 cm^2^, and they had to decompose beyond visible to the naked eye within two days for the compost mixes to qualify for PLA composting tests. PLA used for testing was 5×5 cm^2^, 20 μm thick films (commercially produced by AGT Inc. using raw material from NatureWorks), and unless otherwise noted, they were pretreated in a 65°C water bath for one day to accelerate degradation for the ease of observation. The PLA films were then placed about 5 cm deep into the composts with a thin nylon mesh laid on top of the PLA to facilitate observation. Degradation status of all materials were documented at 5, 10, and 15 days after the start of the composting treatment.

The PLA films used for metagenomics analysis of PLA-associated microbiomes were from three different commercial sources (AGT with raw material from NatureWorks, Jiasheng, Nanochem). For each film, 0.3 g PLA was dissolved in 10 ml dichloromethane (DCM) overnight and poured into a 9-cm glass dish before DCM evaporation in a chemical hood for a second overnight; the resulting films were 25 μm in thickness. The films were subsequently placed into two trained compost bins (S and S3) and one untrained compost (U) for 21 days at which two PLA samples each were collected for trained bins S and S3 and one from the untrained compost. The names of each sample included the compost origin (S/S3/U) as well as the PLA source (A for AGT, J for Jiasheng, N for Nanochem); for example, the PLA film from AGT collected from trained compost bin S was designated as “SA”.

### Scanning electron microscopy

PLA film pieces treated in composts for 15 days were collected, cleaned with ddH_2_O, air-dried, secured onto the sample holders using carbon tape, and coated in gold (Hitachi E-1010 Ion Sputter) for scanning electron microscopy (SEM) imaging (6000X magnification, JEOL JSM-7500F).

### DNA extraction for metagenomics analysis

DNA extraction was harvested from as much leftover PLA sample as possible for collection or 5 g of compost that was not in contact with PLA films. The materials were then placed into 50 ml centrifuge tubes with 10 ml of no carbon media (26.1 mM Na_2_HPO4, 22 mM KH_2_PO4, 8.6 mM NaCl, 18.7 mM NH_4_Cl, 0.4 mM MgSO_4_, 36 M FeSO_4_, 29.6 M MnSO_4_, 47 M ZnCl_2_, 4.6 M CaCl_2_, 1.3M CoSO_4_, 1.4M CuSO_4_, 0.1 mM H_3_BO_3_, 34.2M EDTA, 1.8 mM HCl), vortexed for 15 minutes, and sonicated in an ice water bath for 15 mins before filtration through 6 μm filter papers (ADVANTEC). DNA was subsequently extracted with the DNeasy PowerWater Sterivex Kit (Qiagen) and converted to DNA libraries for whole genome sequencing with the NEXTFLEX Rapid XP V2 DNA-Seq Kit for Illumina Platforms (Revvity). Sequencing of the DNA libraries was performed with the Illumina NovaSeq 150 PE platform commercially (Genomics).

### Data analysis

Raw sequencing reads were first subjected to host genome removal using Bowtie2 [15] human genome reference GRCh38) and assembly with MEGAHIT [16] Next, Prodigal [17] and CD-HIT[18] were used to perform gene predictions and removal of redundant genes before DIAMOND+MEGAN [19] were utilized to obtain taxonomic and enzymatic information in which assembled contigs were blasted against the NCBI non-redundant protein sequence database. Signal peptide predictions were done by SignalP 6.0[20] which assigned five types of signal peptides (Sec/SPI, Sec/SPII, Sec/SPIII, Tat/SpI, Tat/SpII), enzyme categorizations were performed in MEGAN[21] using the Enzyme Commission nomenclature, and Shannon diversity indices were calculated also in MEGAN using PCA.

To determine enrichment levels of specific classes of enzymes, the numbers of proteins in each sample that contained signal peptides belonging to different enzyme categories of the EC nomenclature system were first tallied. Subsequently, these numbers were divided by the total numbers of proteins within an individual sample to determine the relative abundances of signal peptide-containing proteins of each subclass followed by the addition of 0.001 to each value for further calculations. To compare which enzyme subclass was the most enriched on PLA surfaces regardless of their compost bin origin, the relative abundance values for each subclass in each PLA-associated sample was divided by the compost control value (SA/SC, SJ/SC, S3N/S3C, S3J/S3C, UN/UC), and averages of each subclass was then calculated to determine the top 30 subclasses with the highest enrichment levels. To compare enzyme subclass abundances between PLA-associated samples from trained and untrained composts, relative abundance values for each subclass in each PLA-associated sample from trained bins was divided by the corresponding values from the untrained sample (SA/UN, SJ/UN, S3N/UN, S3J/UN). The averages of these four values were then used to identify the subclasses with the highest enrichment.

### PCR and RT-PCR

Microbial filtrates associated with PLA samples from composts were collected as described for extracting DNA for metagenomics analysis. The filtrates were then subjected to DNA and RNA extraction using the EasyPure Genomics DNA Spin Kit (Bioman) and RNeasy PowerMax Soil Pro Kits (Qiagen), respectively. RNA samples were then converted to cDNA using SuperScript IV (ThermoFisher) with random hexamer-priming. To detect gene sequences of three mannonate dehydratases (C1, PQ724380; D1, PQ724381; D3, PQ724382; Fig. S4), nested PCRs were carried out with each round of the reactions consisting of 35 cycles of amplification. The products from the first PCR round were diluted 100-fold before being used as templates in the second round of amplification. The genes and primers used are listed in Table S1.

## Results and discussion

### Mesophilic composts with PLA biodegradability

PLA has been well documented to be compostable under industrial settings in which internal reaction temperatures are higher than 55°C, but the biodegradation has been reported to be minimal when the compost is mesophilic which is typical for composts smaller in volume. We have overcome this limit and developed mesophilic composts with clear PLA biodegradability by continuous culture with PLA material. The mesophilic composts were about 25 L in volume and placed outdoors without any heating elements, thus their internal temperatures ranged from 0-10°C above ambient temperatures which translated to no warmer than 40°C at any time. This activity was transferrable as we have been able to use a pair of bins that first developed mesophilic PLA biodegradability to seed new bins acquire PLA mesophilic compostability.

To demonstrate and document PLA biodegradability of our composts, we placed into the composts thin 100% PLA films that have been pretreated by heat at 65°C for 24 h, a step that accelerated their biodegradation for ease of observation (Fig. S1). In composts trained for PLA biodegradation, the films were rapidly decomposed and were not observable by eye anymore within 10 days whereas PLA films placed into untrained composts remained relatively intact after the same amount of time (samples from untrained bin: UC, samples from trained bins: S and S3, Fig. 1, S2). The composts used for monitoring PLA film degradation were handled minimally during the observation periods to avoid breakage of the pieces due to physical forces. Additional abiotic factors that could affect PLA film integrity such as UV from sunlight and rain were also minimized by placing the composts in lidded bins. However, as insects and worms were inevitably present in the mixtures, we were not surprised to observe some deterioration of the films incubated in untrained composts (Fig. 1A-C), and this impact was expected to be similar in untrained and trained composts. In comparison, the films in trained bins were much more fragmented (Fig. 1E-G, I-K).

**Figure 1.**
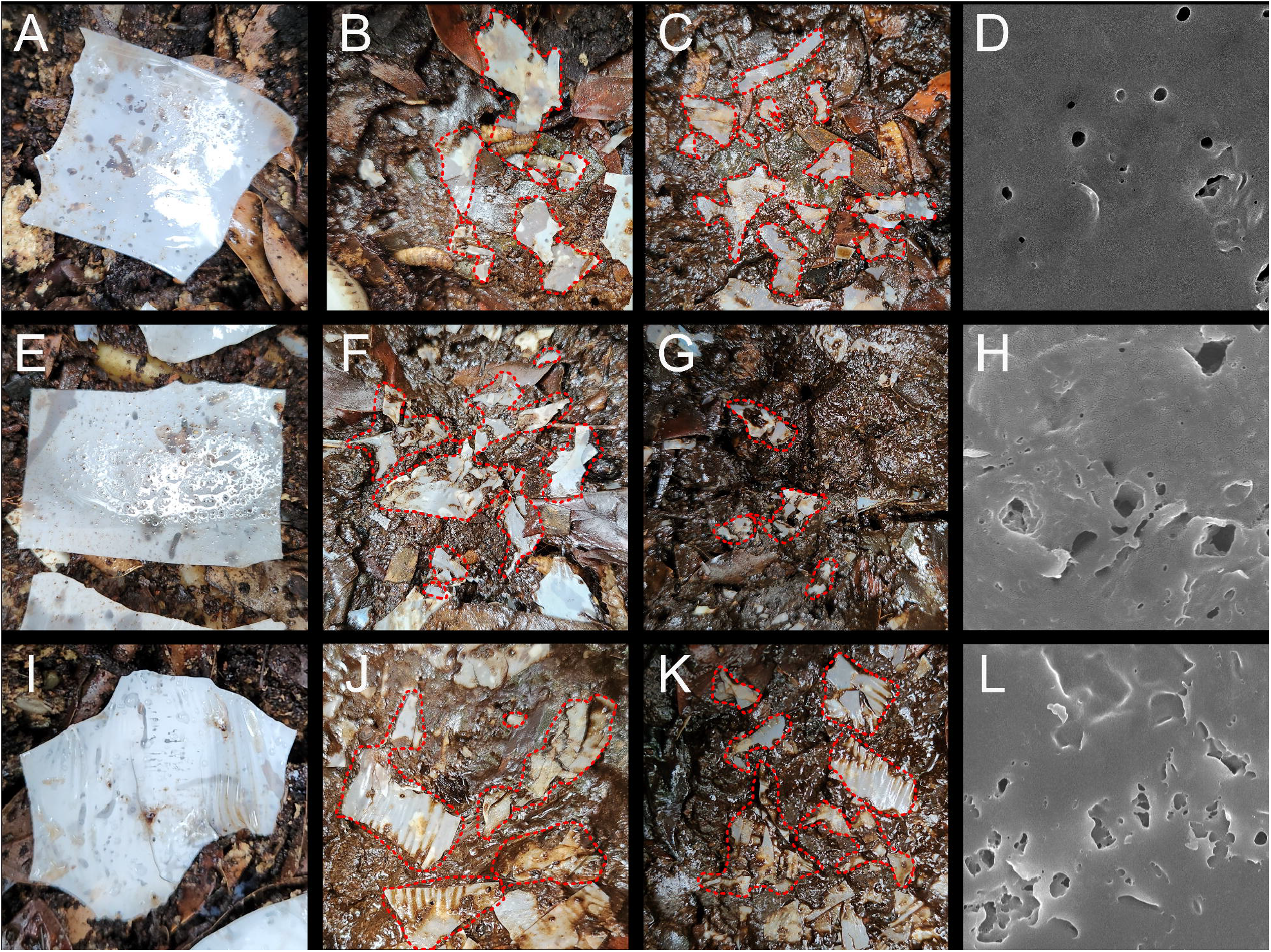
Our trained composts exhibited PLA degradability at mesophilic temperatures. **A-C**, Day 0, 5, and 15 day images of pretreated PLA films placed in an untrained compost. **E-G**, Day 0, 5, and 15 day images of pretreated PLA films placed in the trained bin S. **I-K**, Day 0, 5, and 15 day images of pretreated PLA films placed in the trained bin S3. **D, H, L**, SEM images at 6000X magnification of PLA films after 15 days of compost treatment in untrained compost **(D)**, trained bin S **(H)**, or trained bin S3 **(L)**. The red dotted lines outline the remainder of PLA films.

The films were further imaged by SEM to observe the surfaces at much higher resolution (Fig. 1D, H, L), and we found PLA films incubated in trained composts to have more uneven and digested surfaces (Fig. 1H, L), sometimes exhibiting multiple layers of laminations, and these appearances were absent from films placed into the untrained compost as well as pretreated films that were never placed into composts (Fig. 1D, Fig. S2).

To investigate whether our trained composts contained not only the ability to enhance mesophilic PLA biodegradability, but more importantly, to initiate such processes, we took PLA films that have not been heat-pretreated and placed them in trained and untrained composts. After 15 days, the extents of deterioration observable at the macro level were less obvious (Fig. 2). However, SEM imaging of the surfaces revealed that the films from trained composts were indeed substantially more digested than those from untrained bins (Fig. 2H, L in trained bins vs. Fig. 2D in the untrained bin). These observations indicated that our trained mesophilic composts contained clear PLA biodegradability, and our composts likely contained two types of microbial activity, one that could initiate biodegradation of PLA, and another being the depolymerization of PLA, with both of them being active at lower temperatures.

**Figure 2.**
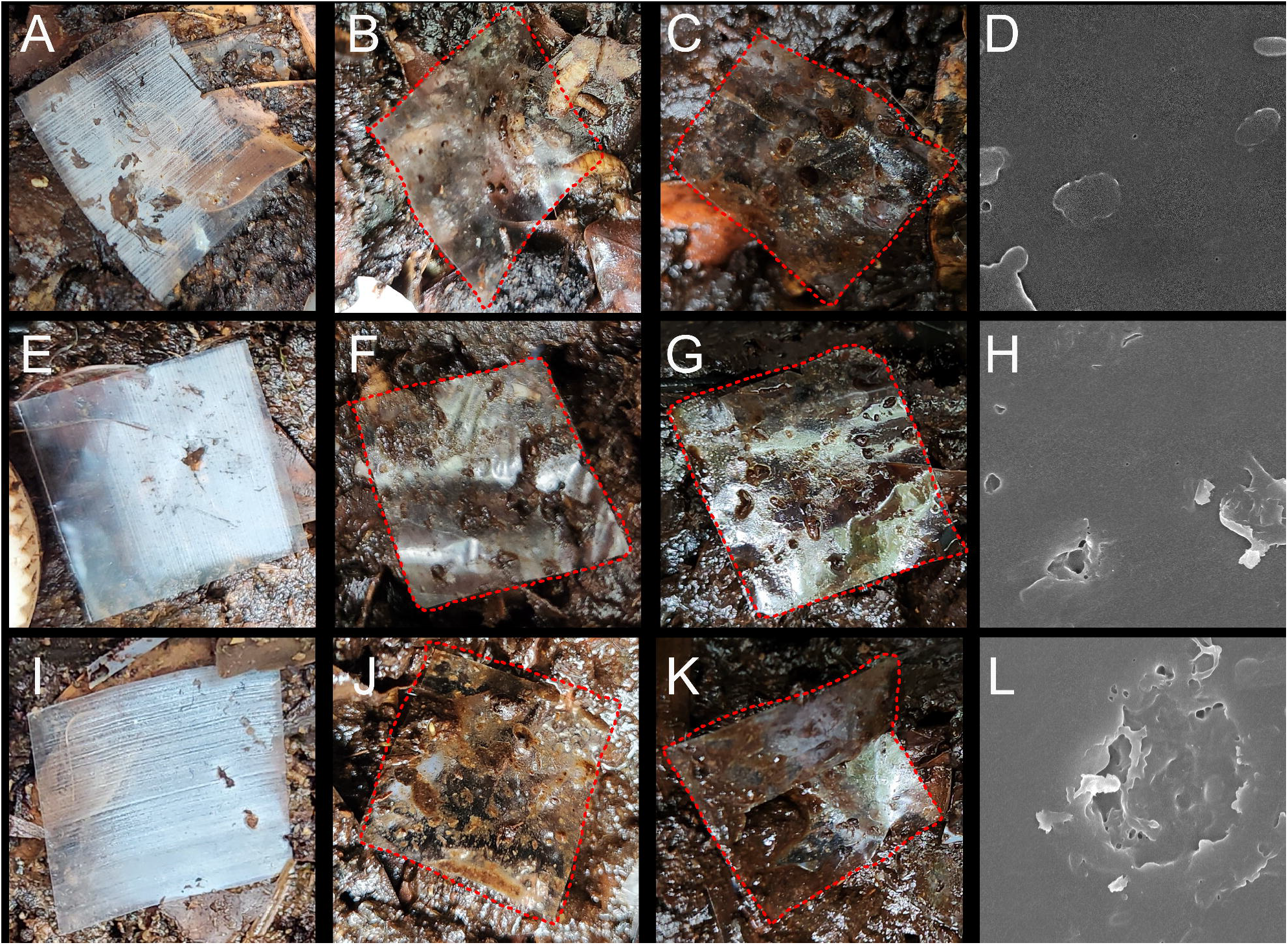
Our trained composts could initiate PLA biodegradation at mesophilic temperatures. **A-C,** Day 0, 5, and 15 day images of untreated PLA films placed in an untrained compost. **E-G**, Day 0, 5, and 15 day images of untreated PLA films placed in the trained bin S. **I-K**, Day 0, 5, and 15 day images of untreated PLA films placed in the trained bin S3. **D, H, L**, SEM images at 6000X magnification of PLA films after 15 days of compost treatment in untrained compost **(D)**, trained bin S **(H)**, or trained bin S3 **(L)**. The red dotted lines outline the PLA films.

### Microbiomes associated with PLA were distinct and were enriched for secreted proteins

Our PLA-degrading mesophilic composts were a valuable resource for studying how PLA breakdown at lower temperatures could be achieved, thus we conducted metagenomics analysis of our trained composts. Specifically, from two trained composts (bins S and S3), we surveyed microbiomes of the surfaces of two PLA pieces each that were undergoing degradation (SA and SN from bin S, S3J and S3N from bin S3, Fig. S1); from these two trained bins, compost material not in touch with PLA was also sampled (SC from bin S and S3C from bin S3, Fig. S1). For comparisons with untrained composts, PLA films were also placed into an untrained compost (U) for 15 days before being harvested for surface microbiome (UN) along with compost material not touching PLA (UC). All trained and untrained composts were fed and maintained with the same green waste for more than 6 months when we sampled them for microbiome profiling, thus the main differences in their compositions would likely be due to their differences in PLA biodegradability.

With the metagenomics data, we first investigated whether they were different based on bacterial species composition using PCA and noticed several trends (Fig. 3A). First, at the species level, samples coming from the same bin were closer to one another than to those from other bins. Second, bacterial composition of samples from the trained bins resembled one another more than to those from the untrained bins. Third, the microbiomes on PLA surfaces from the same bin were more similar to one another than to the PLA-unassociated samples. These pointed to the presence of specific strains related to PLA biodegradation. This point could also be illustrated by comparing the distribution of bacterial orders between samples and their Shannon indices (Fig. S3A-B). Interestingly, when we examined the protein-coding potential of the different samples by PCA, those associated with PLA were clustered together whereas those not in contact with PLA were distinct (Fig. 3B). This again indicated the existence of specific microbial activity associated with PLA compostability.

**Figure 3.**
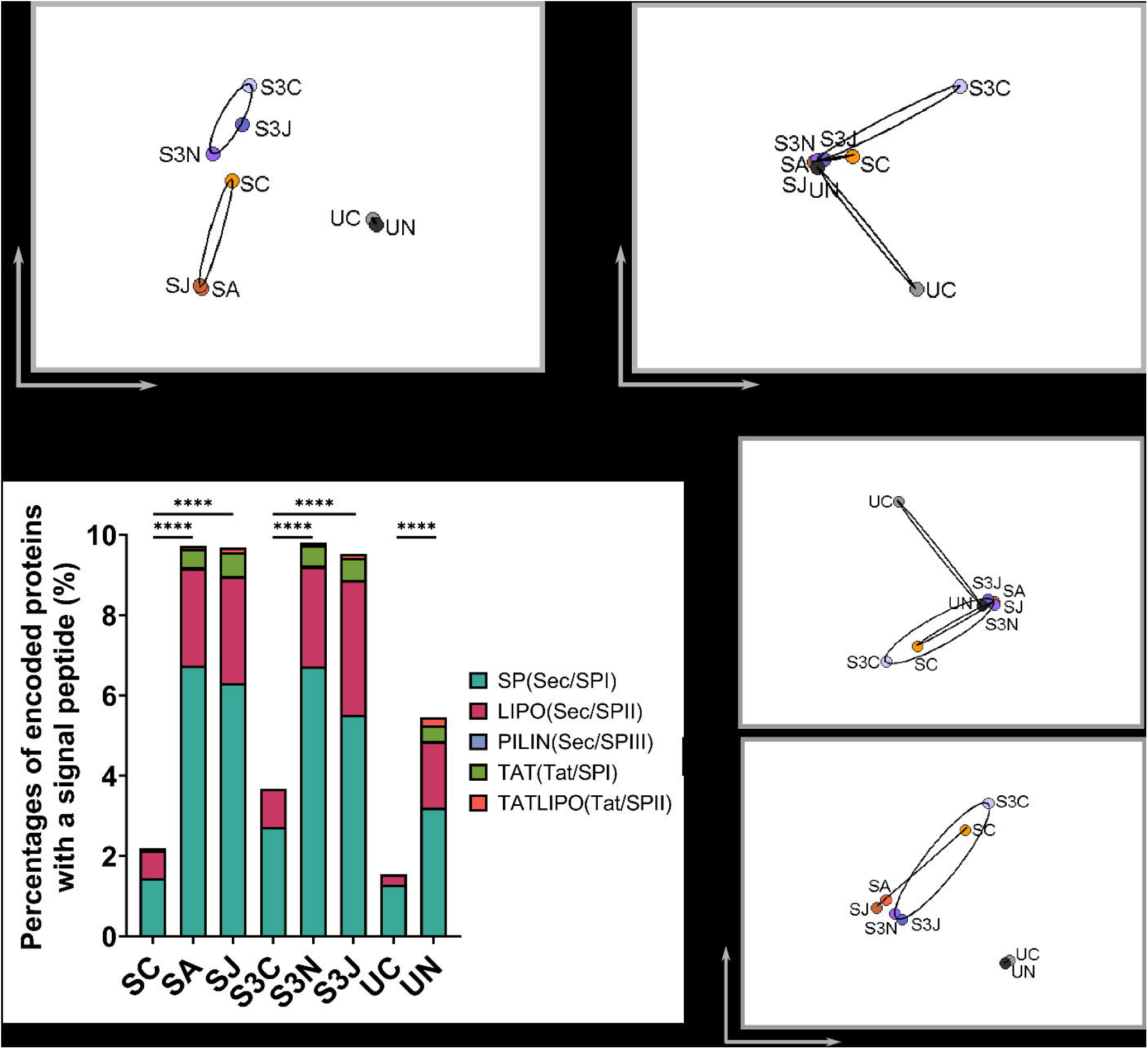
Microbiomes associated with biodegrading PLA were distinct from control materials. **A,** Clustering analysis of all eight samples profiled for their metagenomes using their species composition. The x-and y-axes display the two dimensions that exhibited the highest variations after data transformation. **B,** Comparisons of protein coding potential of the eight metagenomes using PCA. The x-and y-axes display the two dimensions that exhibited the highest variations after data transformation. **C,** Percentages of the eight metagenomes that encoded proteins with any of five canonical signal peptides. ****, p<0.0001. Designations of signal peptide class are displayed on the right. **D,** Clustering analysis of the eight samples based on the composition of species to which signal peptides encoded in each sample originated from. **E,** Clustering analysis of the eight samples based on the consortia of signal peptide-containing enzymes encoded by each metagenome.

PLA polymers are long and their surfaces hydrophobic, thus initiation of biodegradation likely requires secreted enzymes, therefore we performed signal peptide predictions for all proteins determined from the metagenomes. Interestingly, we found the percentages of signal peptide-containing proteins encoded by the PLA-associated metagenomes to be significantly higher than those for unassociated samples for both trained and untrained composts (Fig. 3C). This result again corroborated the hypothesis that there was selection of bacterial strains in the composts for ones that can better colonize on the surfaces of PLA. In addition, the greater potential of strains associated with PLA in producing secreted factors is consistent with the possibility that some of the secreted proteins would be involved in PLA biodegradation.

We next investigated whether there were functional biases of secreted proteins in PLA-associated metagenomes by categorizing them based on EC nomenclature. This revealed that hydrolases and lyases made up higher fractions of all signal peptide-containing proteins encoded in the PLA-associated metagenomes compared to unassociated samples (Fig. S3C). These differences were not present when all proteins encoded by the metagenomes were used for analysis (Fig. S3D), indicating that the higher occurrences of hydrolases and lyases were specific to secreted proteins and suggested that this phenomenon could be related to PLA biodegradability. This subsequently prompted us to carry out PCA-based clustering analysis of the eight samples using only the signal peptide-containing proteins they encoded. We found that if this analysis was done using the enzyme subclasses represented in each sample, the PLA-associated samples were closely clustered together and separated from those unassociated with PLA (Fig. 3D). If clustering was carried out using the bacterial species from which the signal peptide-containing proteins originated, the eight samples became grouped into three clusters (Fig. 3E): PLA samples from trained composts (SA, SJ, S3N, S3J), control samples from trained composts (SC, S3C), and both samples from the untrained compost (UC, UN). This suggested that there were clearly strain differences between the trained and untrained composts. Moreover, the strains that encode signal peptide-containing proteins are quite similar among the two trained composts. Taken together, these analysis results point to specific strains in the trained composts producing secreted factors to mediate their PLA degradability.

### Serine hydrolases were enriched in the genomes of PLA-associated microbes

To use our metagenomics data to identify specific enzymes that could mediate PLA biodegradation, we performed clustering analysis with separate enzyme classes for signal peptide-containing proteins and found hydrolases and lyases with patterns consistent with them being involved in PLA biodegradability (Fig. S4). For hydrolases, all PLA-associated samples were clustered together away from the unassociated ones (Fig. S4C). In the case of lyases, the PLA-associated samples from trained composts were clustered near each other but away from that from untrained compost (UN) and also away from the samples unassociated with PLA (Fig. S4D).

A different approach for finding enzymes of interest was to calculate and rank the enrichment levels of individual enzyme subclasses between PLA-associated and unassociated samples. This was also performed with signal peptide-containing proteins belonging to individual EC enzyme subclasses. The enzyme subclass with the highest enrichment level based on this calculation was serine-type D-Ala D-Ala carboxypeptidase (EC 3.4.16.4), a type of serine protease (Fig. 4A). In addition to this subclass, many other hydrolase subclasses also showed strong enrichment levels (Fig. 4A), which is consistent with the general class of hydrolases (EC 3) being over-represented in PLA-associated samples (Fig. S4C). The finding of a serine protease at the top of the enrichment list comparing PLA-associated and unassociated samples reaffirmed results from previous studies reporting serine proteases to be the main type of enzyme with PLA depolymerization activity.

**Figure 4.**
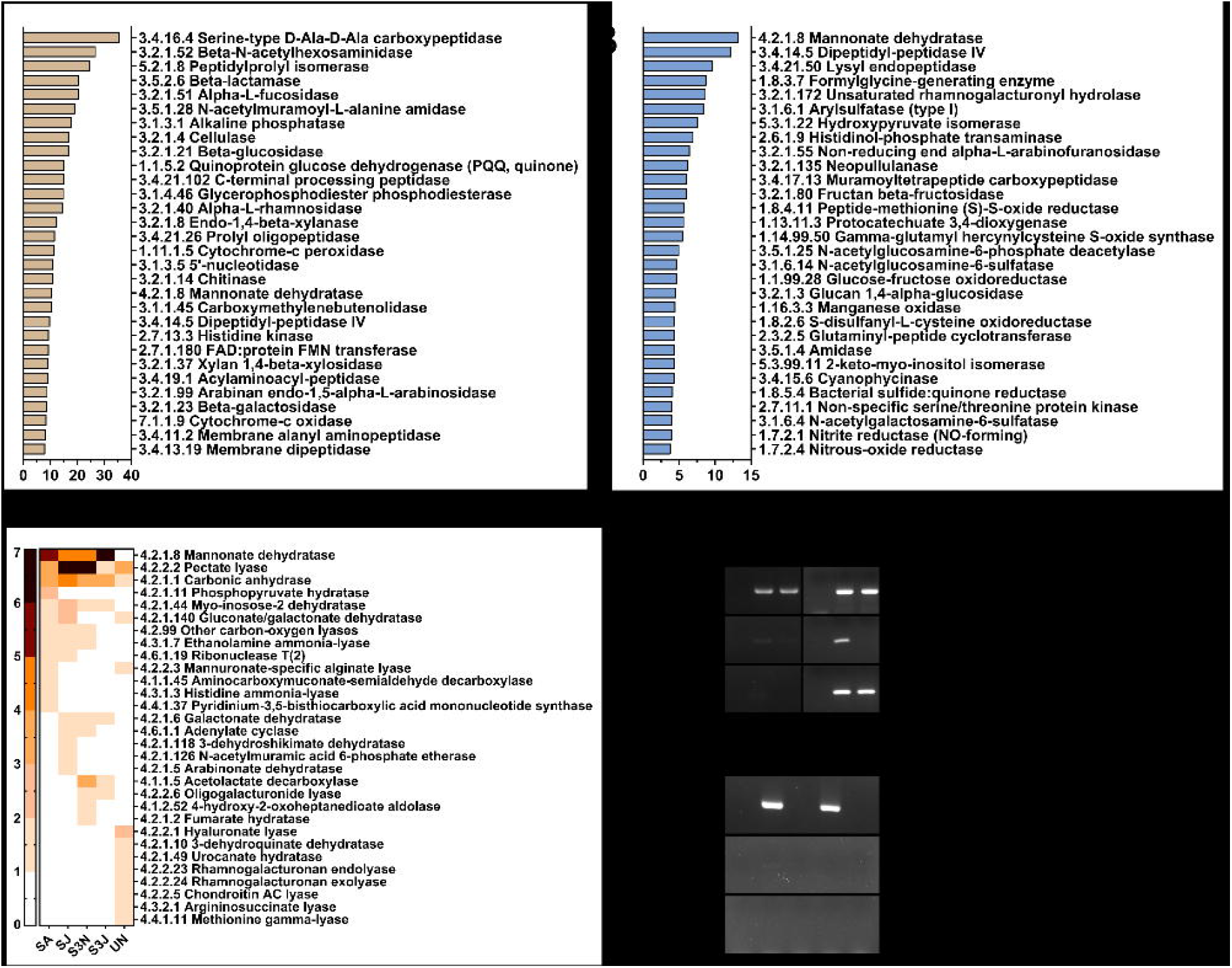
Enzyme subclasses that exhibited enrichment on the surfaces of PLA undergoing biodegradation. **A,** Enzyme subclasses most over-represented comparing all PLA-associated samples (SA, SJ, S3N, S3J, UN) to those that were not (SC, S3C, UC). **B,** Enzyme subclasses most over-represented comparing PLA-associated samples from trained composts (SA, SJ, S3N, S3J) to that from the untrained compost (UN). **C,** Occurrences of subclasses of hydro-lyases among signal peptide-containing proteins encoded by the metagenomes associated with PLA. Enzyme subclasses were ordered by decreasing average occurrences in the four PLA-associated samples from trained composts (SA, SJ, S3N, S3J), and the darkness of colors indicate higher occurrences of the indicated subtypes in individual samples. **D,** Nested PCR amplification of three mannonate dehydratase encoding-genes, C1, D1, and D3 (Fig. S4). One of the DNA templates came from PLA harvested from the trained bin S whereas the other contained PLA from three trained bins, S, S2 (also related to S), and S3. For both **(D)** and **(E)**, two annealing temperatures for the second round of PCR amplification were used, 52°C and 54°C, and they are indicated on the panels. NTC indicates the no-template control. **E,** RT-PCR amplification of the three mannonate dehydratase under the same nested PCR protocol as in **(D)** using RNA from PLA surfaces collected from the trained bin S. There were strong signals for C1 and a faint band for D1 with 52°C as the annealing temperature. +RT and -RT refer to the reverse transcription reactions done with or without addition of reverse transcriptase.

### Hydro-lyase was the most strongly-enriched enzyme subclass on biodegrading PLA surfaces

A key activity we wanted to investigate was the one that would distinguish our mesophilic trained composts from untrained composts. For this, we calculated and ranked enrichment levels of enzyme subclasses comparing PLA-associated samples from trained bins to that from the untrained bin, and for this analysis we also used signal peptide-proteins predicted for each sample. This list was topped by mannonate dehydratase (EC 4.2.1.8, Fig. 4B), a subclass of lyases that catalyzes breakage of carbon-oxygen bonds. This was very intriguing as lyases was the only major enzyme category for which clustering analysis resulted in separation between the PLA-associated samples from trained composts and the untrained one (Fig. S4D), thus we further analyzed the origin of this separation by examining the occurrences of all subclasses of lyases in all PLA-associated samples (Fig. 4C). There were two notable observations from this analysis. First, the abundance of lyases were higher in PLA-associated samples from trained composts than from untrained composts. Second, the subclasses of lyases found in PLA-associated samples from trained vs. untrained composts were different. Mannonate dehydratase was the most abundant type of lyase encoded by PLA-associated microbes of trained composts while being completely absent in controls (SA/SJ/S3N/S3J vs. UN, Fig. 4C).

To validate the importance of mannonate dehydratases in our system (Fig. S5), we collected independent samples from trained composts and tested if three genes encoding mannonate dehydratases that exhibited strong enrichment in biodegrading PLA samples from the metagenomics data could again be detected. Using nested PCR, we were able to detect the presence of all three mannonate dehydratase-encoding genes in trained composts (Fig. 4D). We further investigated whether these enzymes were expressed in the trained composts and were able to detect transcripts for two of the three genes in the trained bin S (Fig. 4E).

These results strongly supported the idea that certain hydro-lyases contribute functionally to the process of PLA biodegradation, and they could be particularly important at mesophilic temperatures. We will note that, in addition to lyases, several proteases also exhibited clear enrichment of trained vs. untrained composts among PLA-associated samples (Fig. 4B), reaffirming the importance of proteases to PLA biodegradation. Taken together, our metagenomics analysis has revealed several enzyme subclasses that are likely to be responsible for PLA biodegradability under mesophilic conditions.

Mannonate dehydratase is a subtype of hydro-lyases whose common activity is to catalyze cleavage of ester bonds without water. As carbon-oxygen bonds are part of the PLA backbone, one possibility is that hydro-lyases is a type of enzyme that can depolymerize PLA in addition to proteases. However, it is tempting to hypothesize that hydro-lyases provide the activity for the initiation step of PLA breakdown. Proteases require water molecules for their cleavage activity which would be difficult on PLA surfaces that are very hydrophobic. In contrast, hydro-lyases, which do not utilize water for their enzymatic activity, could be a candidate for PLA surface functionalization. Such a role would be consistent with multi-step plastic biodegradation models in which surface digestion or hydrolysis occurs before enzymatic depolymerization would take place [4]. For PLA, the first step could be abiotic—such as chemical hydrolysis in thermophilic composts—or biotic, a novel activity we propose our mesophilic compost to harbor. Such a biological activity would be advantageous as the lowering of reaction temperatures could broaden the scenarios for which PLA biodegradation could be applicable, and the energy requirement for these applications would also be reduced, all of which would further increase the feasibility of PLA biodegradation.

## Data availability

All raw and processed sequencing data generated in this study have been submitted to the NCBI BioProject database under the accession number PRJNA1185390.

## Acknowledgements

The authors would like to thank the Yang lab for comments and suggestions. Data analysis was performed at the CGU AI center and the National Center for High-Performance Computing, SEM imaging was done at the Microscopy Center at Chang Gung University. This work was supported by an NSTC grants (112-2311-B-182-003-MY3) to S.Y.Y.

## Author contributions

Conceptualization, S.D.F.,S.Y.Y; data curation, S.W.H., S.D.F., A.C.K.; formal analysis, S.W.H., C.-F. L.; writing, S.Y.Y., S.W.H., T.S.M.A.; funding acquisition, S.Y.Y.

