## Supplementary data for "Mesophilic compostability of polylactic acid and the associated microbiome as revealed by metagenomics"

**Supplementary Information**

**Supplementary Table**

**Table S1.** Primers used for nested PCR of mannonate dehydratase encoding-genes.

| Mannonate dehydratase genes^a^ | PCR round | Forward primer (5’-3’) | Reverse primer (5’-3’) |
| --- | --- | --- | --- |
| C1 | 1 | GGGATTGAGCGGACATGGAG | GCAGCTGGAGGATCATTGGG |
|  | 2 | GCAAACAGGCAACGGATGAG | TCCACAACGGGCAATCCAAT |
| D1 | 1 | GAGGGACTATGTGCGGGATG | TTTGGTGCTCGGTATCACCC |
|  | 2 | AAGCCTTGGTACGCTGACAA | CTGGCACCACTGCTTTCAGA |
| D3 | 1 | GCAGCTGTGGGGGTAGTAAAT | GCCAAATTCGGTCAATGGCA |
|  | 2 | CTTACGGGCCAAAGAGCAGA | AATCACCTCAATACCGGCCC |

^a^Genes names are as designated in Fig. S4.

**Supplementary figures and figure legends**

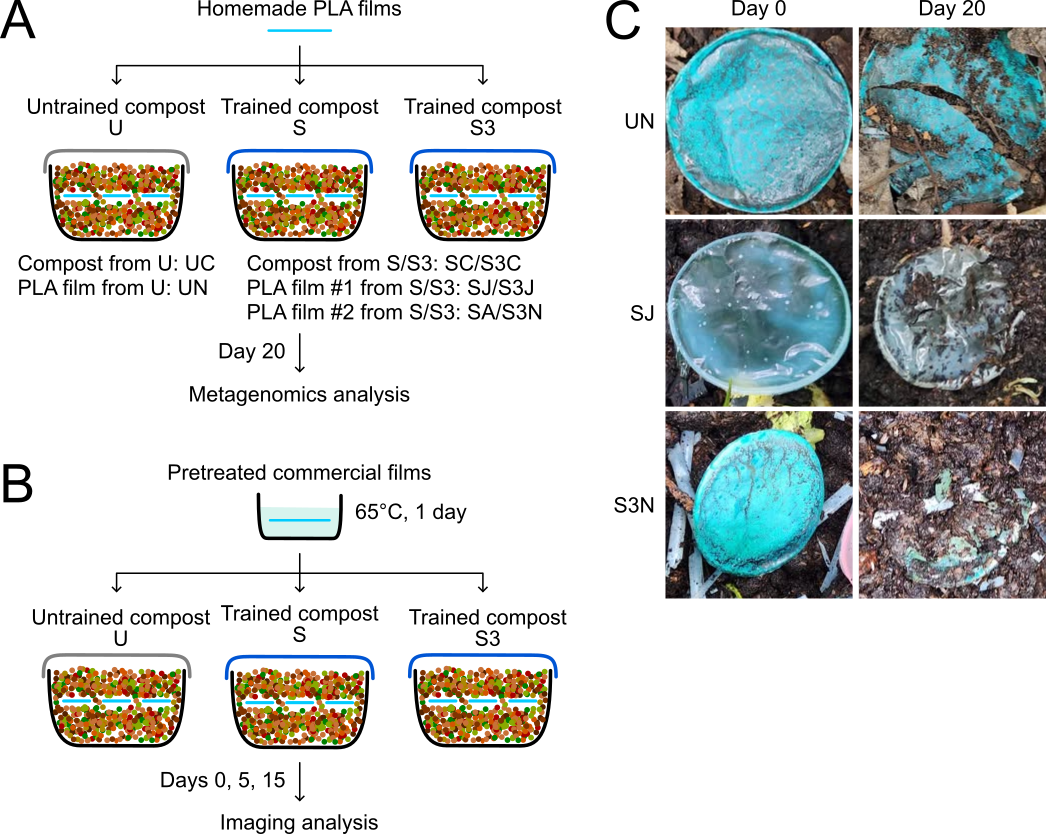

**Figure S1.** Schematic diagrams of our sample collection strategies. **A**, The work flow for how samples for metagenomics profiling were collected. **B**, Diagram of how PLA compostability was tested. **C**, Images of PLA films sampled at day 20 for DNA extraction for metagenomics analysis and at day 0 for comparisons.

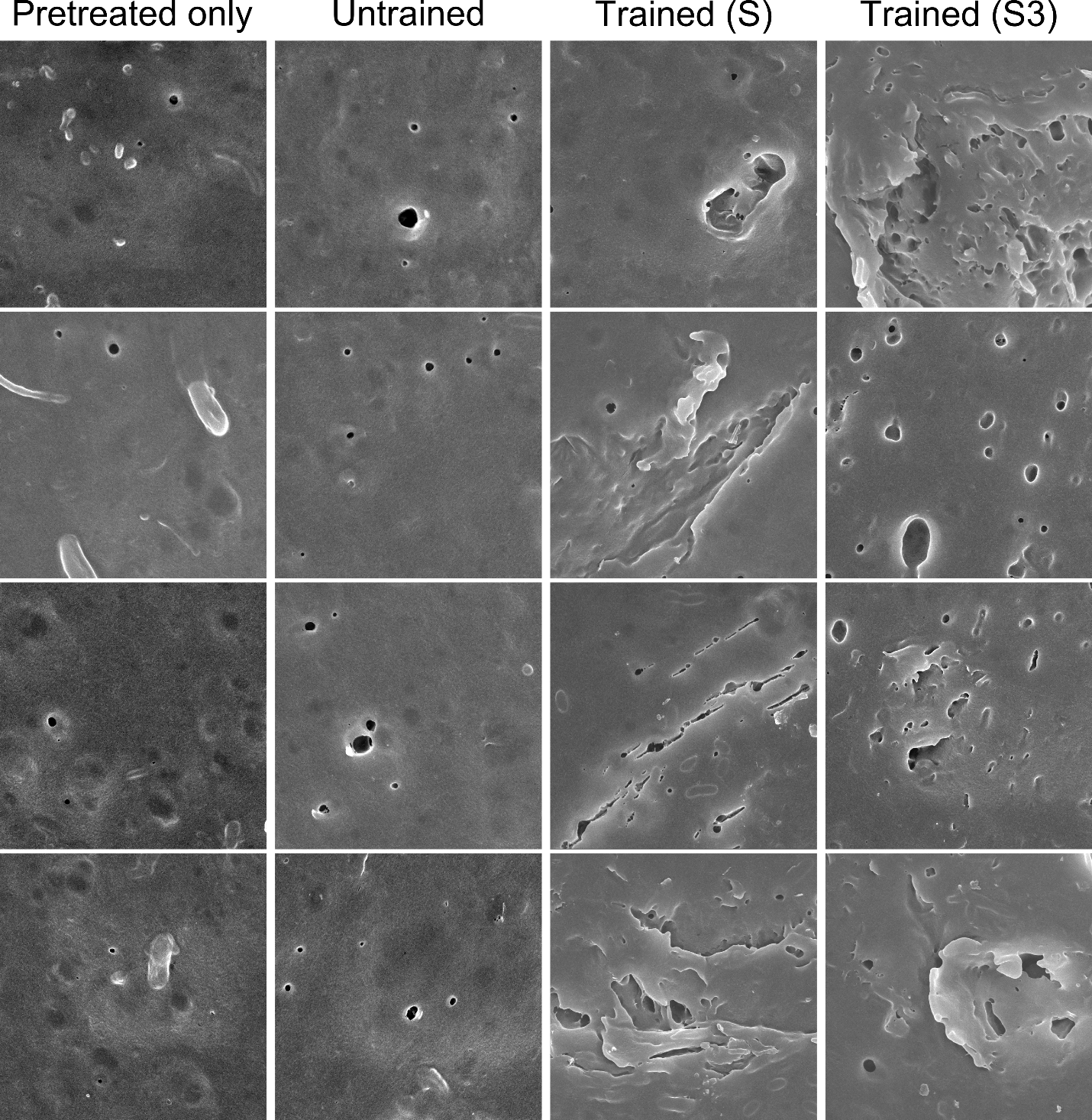

**Figure S2.** Additional SEM images documenting surfaces of PLA films after different compost exposure as indicated. Films that have been pretreated but not placed into composts were also imaged as controls (left column).

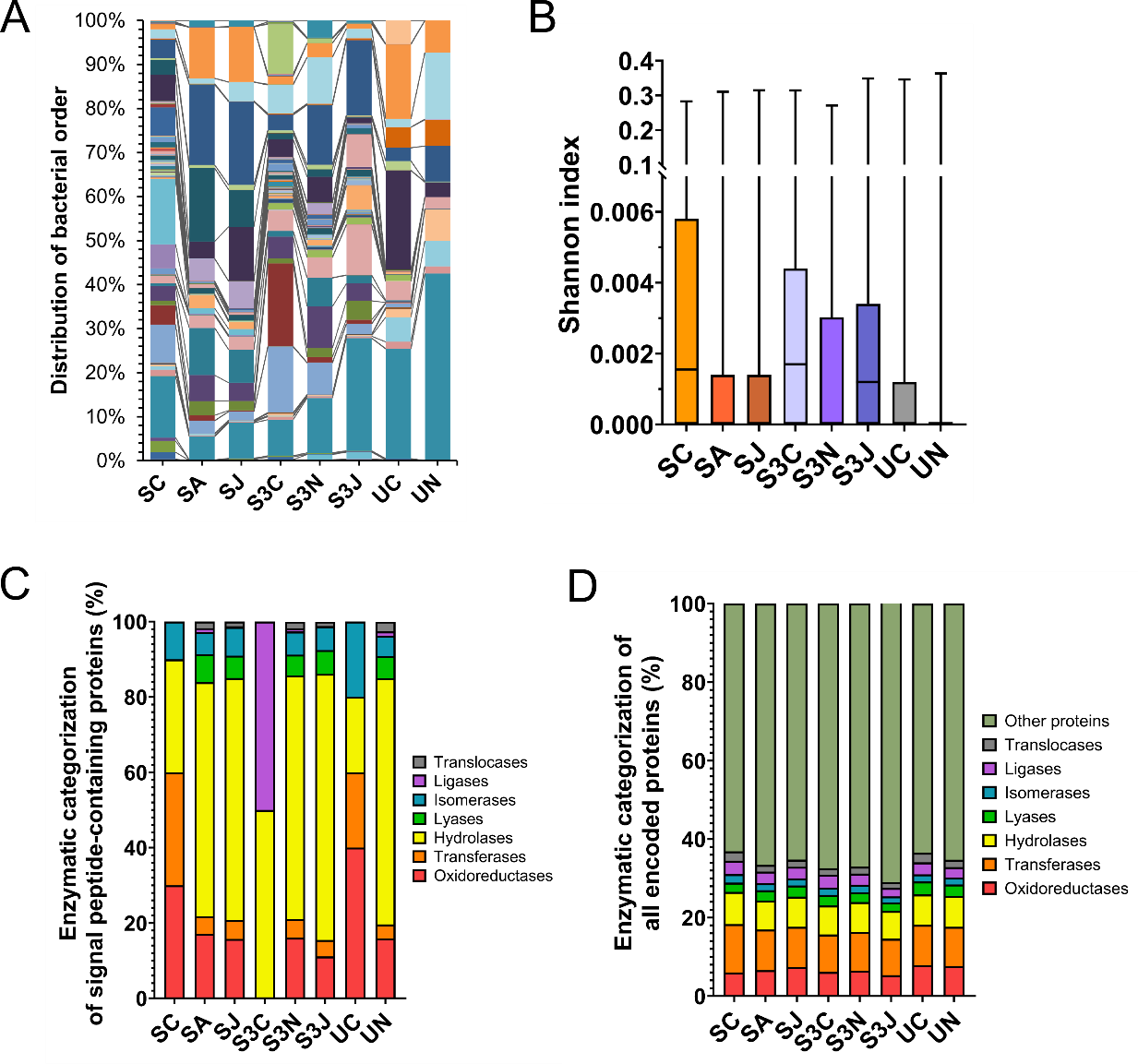

**Figure S3.** Analysis of the microbiome composition of different compost samples. **A**, Differences in bacterial strain distribution at the level of orders between different samples. The lines connecting between bars indicate the same bacterial order. **B**, Shannon indices which reflects intra-sample diversity of all eight samples. **C**, Distribution of signal peptide-containing proteins encoded by each metagenome into functional categories. **D**, Distribution of all proteins encoded by each metagenome into functional categories. For **C**-**D**, names of enzyme classes are listed on the right.

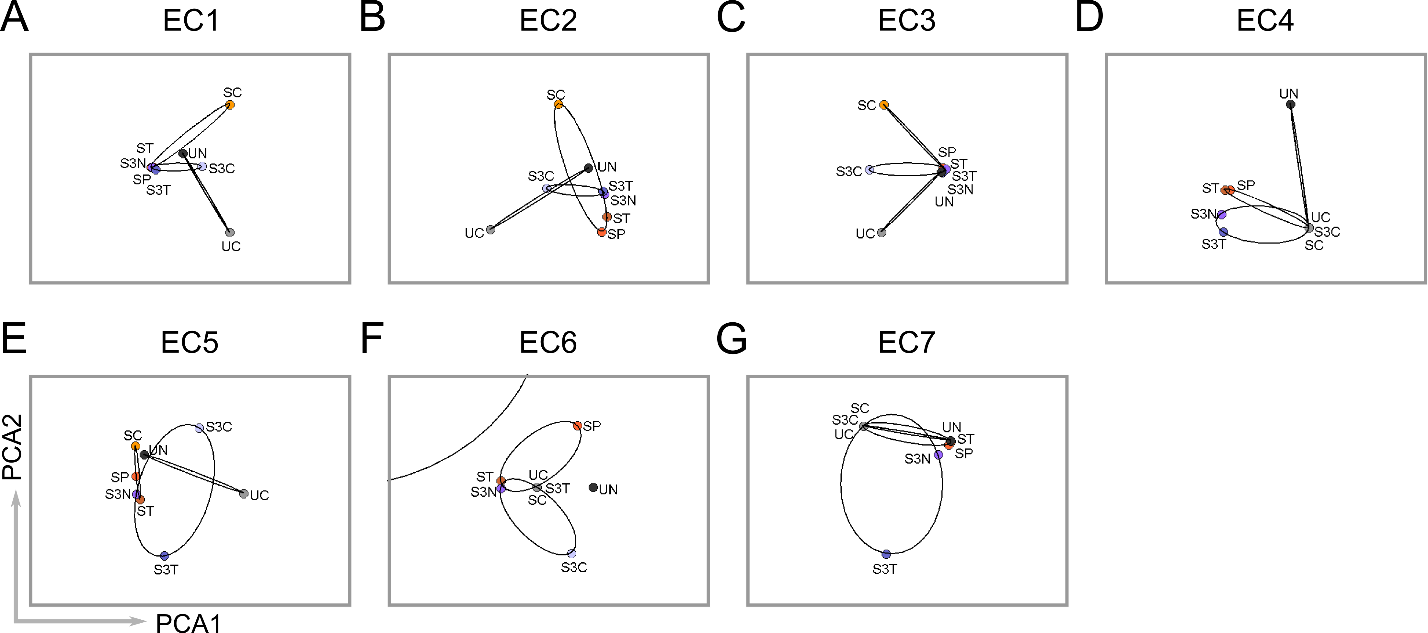

**Figure S4.** PCA-based clustering analysis of functionally categorized signal peptide-containing proteins encoded by the metagenomes of all eight samples. **A**, Oxidoreductases (EC1). **B**, Transferases (EC2). **C**, Hydrolases (EC3). **D**, Lyases (EC4). **E**, Isomerases (EC5). **F**, Ligases (EC6). **G**, Translocases (EC7).

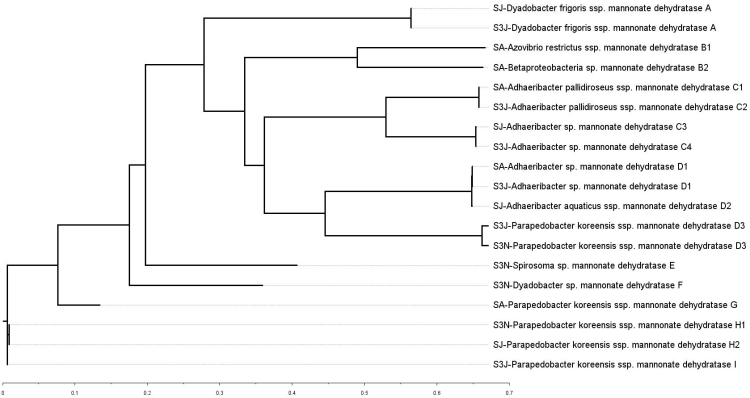

**Figure S5.** Phylogenetic tree of genes encoding mannonate dehydratases detected in the trained composts from our metagenomic profiles (SA, SJ, S3N, S3J). Each entry represents an ORF constructed from separate samples, and the sequences that were 100% identical were given the same names.
